# Frequent extrachromosomal oncogene amplification drives aggressive tumors

**DOI:** 10.1101/859306

**Authors:** Hoon Kim, Nam Nguyen, Kristen Turner, Sihan Wu, Jihe Liu, Viraj Deshpande, Sandeep Namburi, Howard Y. Chang, Christine Beck, Paul Mischel, Vineet Bafna, Roel Verhaak

## Abstract

Extrachromosomal DNA (ecDNA) amplification promotes high oncogene copy number, intratumoral genetic heterogeneity, and accelerated tumor evolution^1–3^, but its frequency and clinical impact are not well understood. Here we show, using computational analysis of whole-genome sequencing data from 1,979 cancer patients, that ecDNA amplification occurs in at least 26% of human cancers, of a wide variety of histological types, but not in whole blood or normal tissue. We demonstrate a highly significant enrichment for oncogenes on amplified ecDNA and that the most common recurrent oncogene amplifications arise on ecDNA. EcDNA amplifications resulted in higher levels of oncogene transcription compared to copy number matched linear DNA, coupled with enhanced chromatin accessibility. Patients whose tumors have ecDNA-based oncogene amplification showed increase of cell proliferation signature activity, greater likelihood of lymph node spread at initial diagnosis, and significantly shorter survival, even when controlled for tissue type, than do patients whose cancers are not driven by ecDNA-based oncogene amplification. The results presented here demonstrate that ecDNA-based oncogene amplification plays a central role in driving the poor outcome for patients with some of the most aggressive forms of cancers.

---

Somatic gain of function alterations in growth controlling genes, especially driver oncogenes, plays a central role in the development of cancer^4–6^. Oncogene amplification is one of the most common gain of function alterations in cancer, enabling tumor cells to circumvent the checks and balances that are in place during homeostasis and providing selective and autonomous advantage to drive tumor growth. EcDNA-based amplification has long been recognized as a way for cells to increase the copy number of specific genes^7,8^, but their frequency appears to be vastly underestimated^2,9^. EcDNA amplification has recently emerged as a powerful mechanism for enabling tumors to concomitantly reach high copy of growth promoting genes, while still maintaining intratumoral genetic heterogeneity through its non-chromosomal mechanism of inheritance^1–3^. To date, cytogenetic methods requiring live cells in metaphase have been used to infer intranuclear localization of DNA amplifications and extrachromosomal status^10^. Consequently, it has been challenging to accurately assess the frequency, distribution, and clinical impact of ecDNA-based amplification. More recently computational analyses of whole-genome sequencing data have suggested a relatively high frequency of ecDNA in some cancer types^11,12^. Here we set out to perform a global survey of the frequency of ecDNA-based oncogene amplification, while investigating its contents and determining its clinical context.

EcDNA are characterized by two distinguishing properties: 1. ecDNAs are highly and focally amplified and 2. they are circular. These properties provide a basis for the AmpliconArchitect tool, that enables detection and characterization of ecDNA from whole-genome sequencing data (**Fig. 1A**)^11^. We applied AmpliconArchitect^11^ to whole-genome sequencing data from The Cancer Genome Atlas (TCGA), to quantify and characterize the architecture of amplified regions that are larger than 10kb and have more than 4 copies (CN>4) above median sample ploidy (**Supplementary Table 1**). Amplicons were classified as ‘Circular’ (**Extended Data Fig. 1A**) representing amplicons residing extrachromosomally or ecDNA structures that reintegrated into non-native chromosomal locations as homogenously staining regions (HSRs), ‘Amplified-noncircular’ for linear amplifications, or as ‘heavily rearranged’, for non-circular amplicons containing segments from different chromosomes, or regions that were very far apart on chromosomes (>1Mb) regions. Sample lacking amplifications were labeled ‘no copy number amplification (CNA) detected’.

**Fig. 1.**
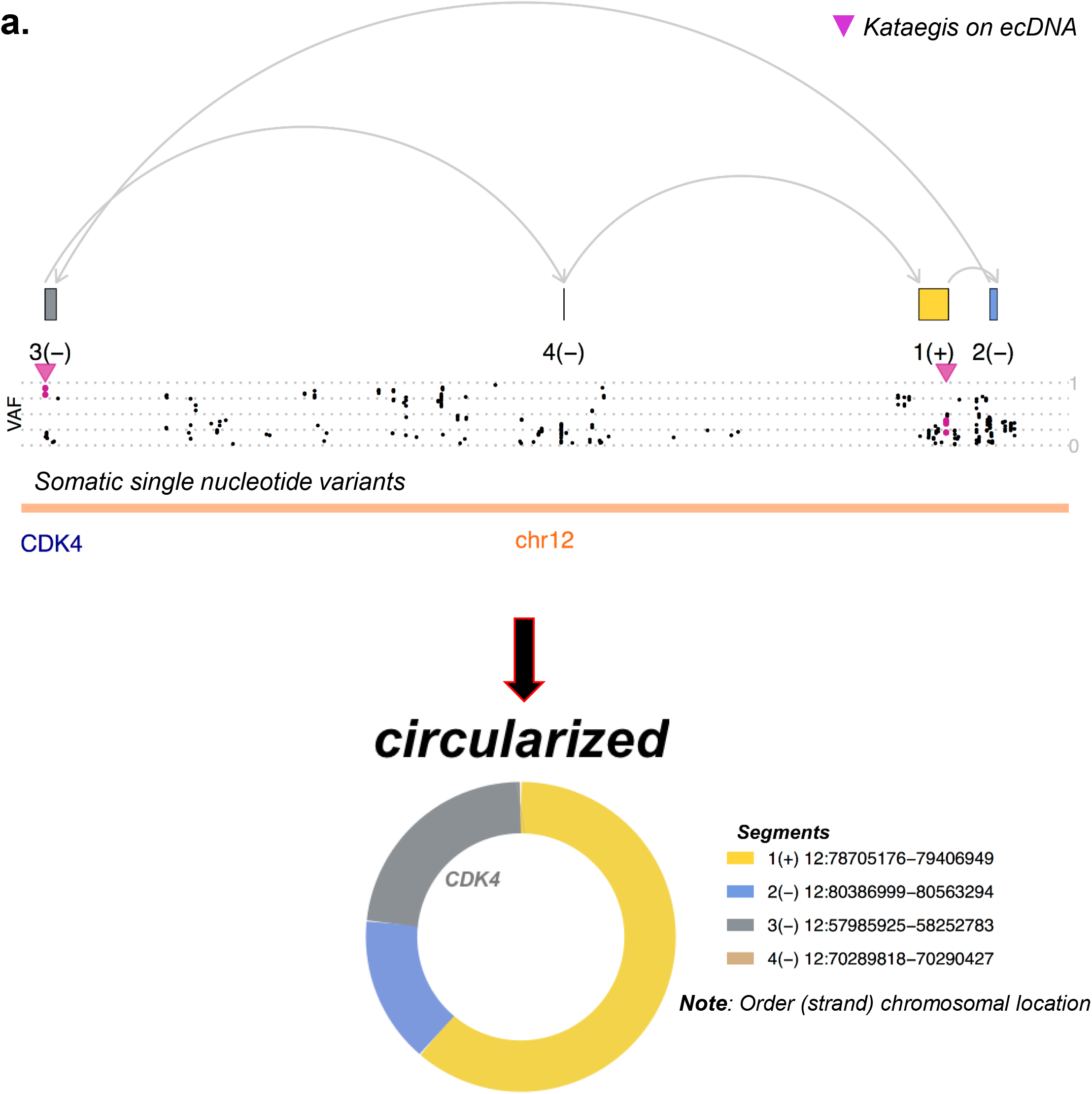
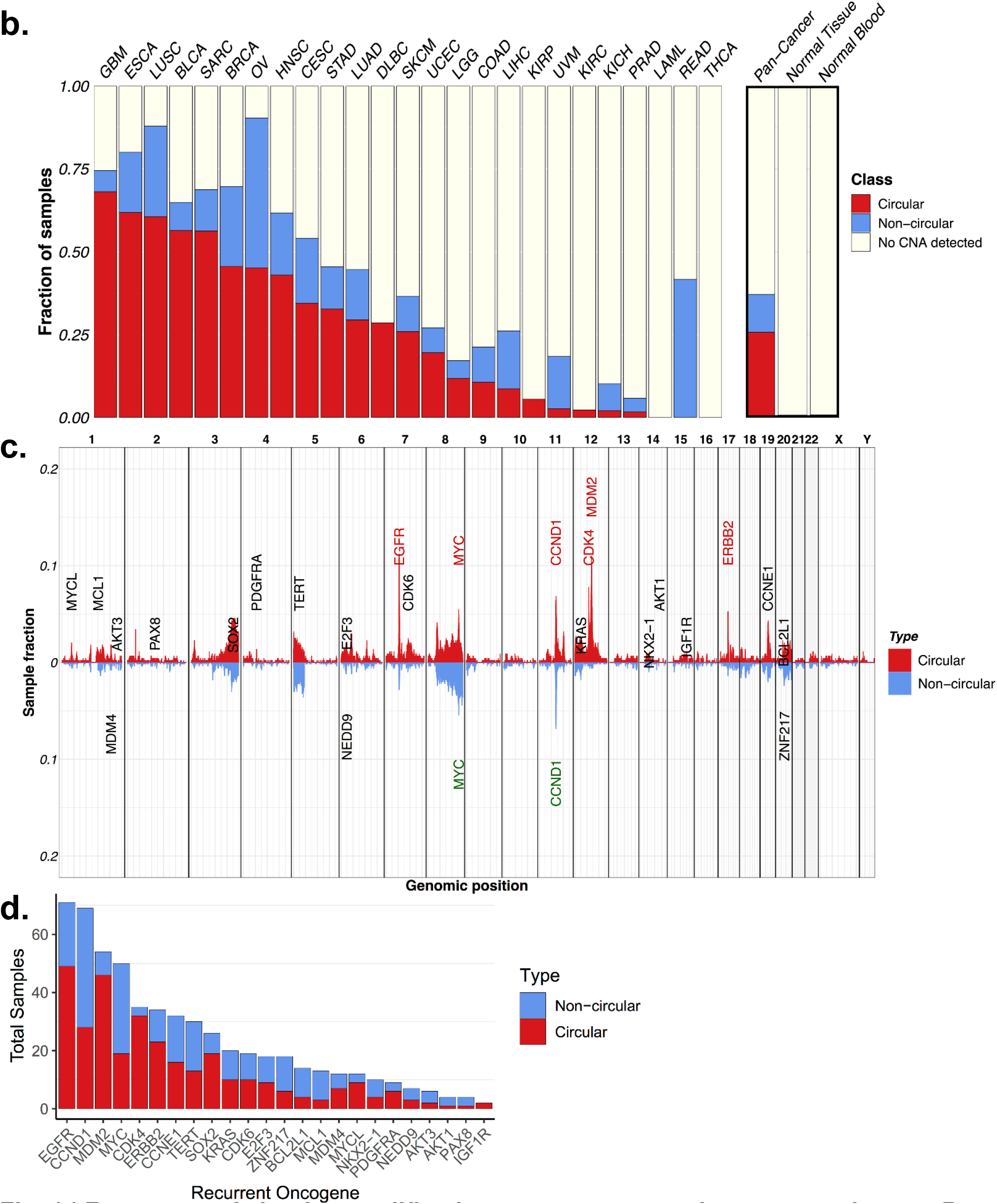
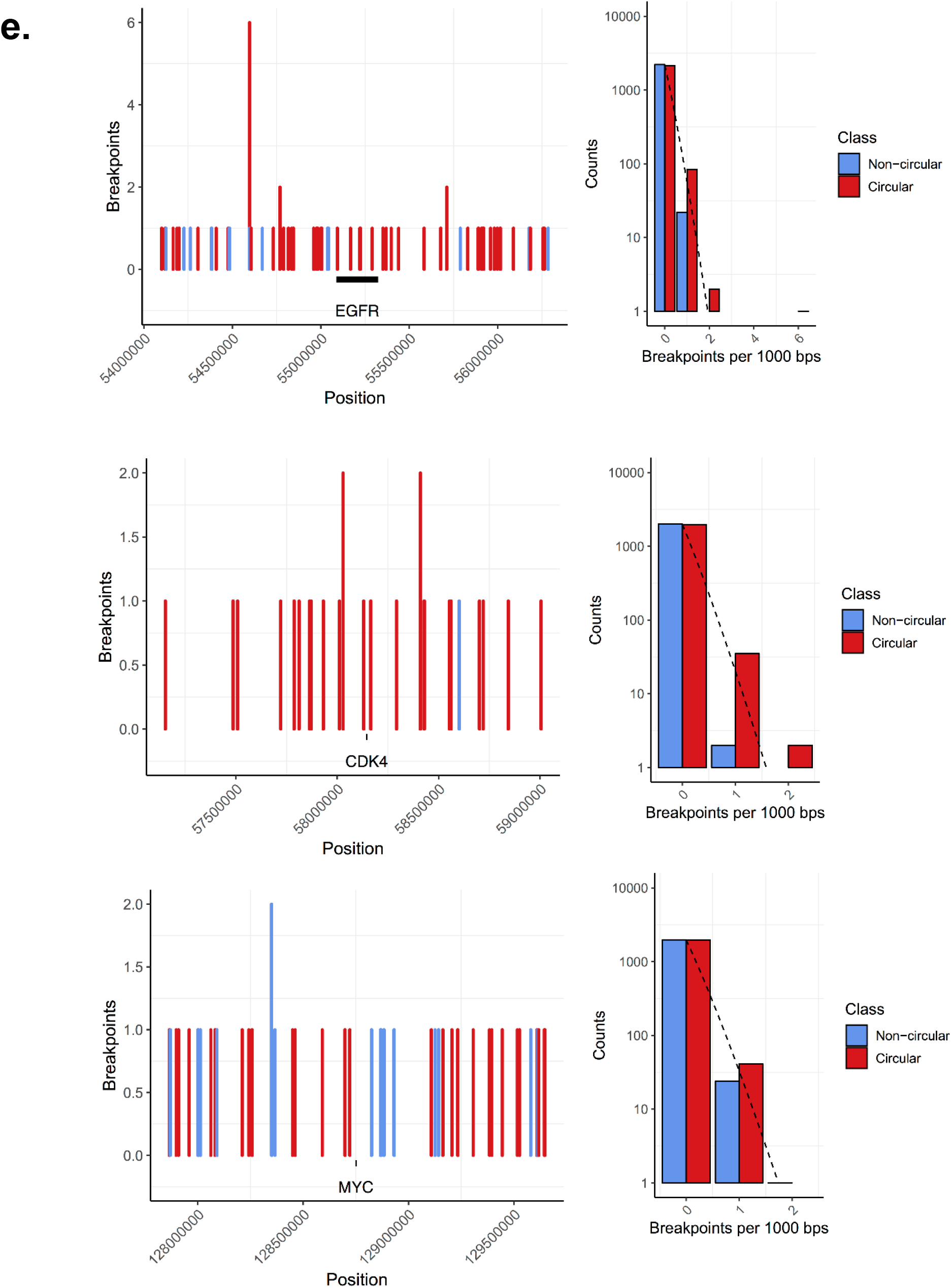
Frequency of circular amplification across tumor and non-tumor tissues. **A**. Representative example of a Circular DNA amplification. **B.** Distribution of circular, non-circular, and no copy number alteration (no CNA) detected categories by tumor and normal tissue. **C**. Genome-wide distribution of circular (red) and non-circular (blue) amplification peaks. **D**. Classification of circular vs non-circular amplification status by gene. Shown are 24 most frequently amplified oncogenes. **E**. Breakpoint locations and distribution of breakpoints across all samples with amplified EGFR (top), CDK4 (middle), and MYC (bottom).

To evaluate the accuracy of the computational predictions, we similarly analyzed whole genome sequencing data from a panel of 34 cancer cell lines^1,2^, for which tumor cells in metaphase could be examined. We used 15 unique fluorescence in-situ hybridization (FISH) probes in combination with matched centromeric probes (60 distinct “cell-line, probe” combinations) to determine the chromosomal or extrachromosomal location of a set of amplicons. We observed that 100% of amplicons characterized as ‘Circular’ by whole genome sequencing profile demonstrated extrachromosomal fluorescent signal (**Extended Data Fig. 1B**). Circular amplicons had a median count of 14.5 ecDNA per cell, in contrast with the ‘Amplified-noncircular’ category, which had a median count of 0.0 ecDNA per cell. However, ecDNAs may be undercounted in amplicons with low copy number. ‘Heavily rearranged amplicons’ showed at least one ecDNA per cell in two of five cases, suggesting that this category consists of a mixture of chromosomal and extrachromosomal amplifications. We excluded the more ambiguous category of ‘heavily rearranged’ amplicons from futher comparisons, confining our analysis to 1,695 TCGA samples. The analytic results of the 256 samples containing the more ambiguous ‘heavily rearranged amplicons’ are presented in the supplement (**Extended Data Fig. 2**).

We found that 436 (26%) of the 1,695 tumor samples carried one or more Circular amplicons, suggesting that ecDNA-based amplification is a common event in human cancer (**Fig. 1B**). In contrast, Circular amplifications were found in <0.5% of matched whole blood or normal tissue samples, suggesting that extrachromosomal amplification is a mechanism that is used primarily by cancer cells (**Fig. 1B**). Of note, our analysis does not reflect the presence of circulating cell free DNA in blood, or of small (<1 kb), circular, non-amplified DNAs, that have been shown to be common in non-neoplastic and tumor tissues ^13–15^. EcDNA-based Circular amplicons were found in all cancer types except acute myeloid leukemia and thryroid carcinoma, including at high frequency in many cancers that are considered to be amongst the most aggressive histological types. The distribution of Circular amplicon frequencies across the samples are consistent with earlier results on cancer models, showing that ecDNA driven amplifications were a defining feature of multiple cancer sub-types, but not normal cells^2^.

The chromosomal distribution of the 627 Circular amplicons was highly non-random (**Fig. 1C**), more so when compared to the Amplified-noncircular regions (**Extended Data Fig. 3A**). We found that 41% of the 24 most recurrent amplified oncogenes were most frequently present on Circular amplicons, with frequencies ranging from 25% of samples for *PAX8* to 91% for *CDK4* (**Fig. 1D**). The result carried over to a larger list of 707 genes that were amplified in at least five samples, with 41% of those oncogenes most frequently being amplified on circular structures (**Extended Data Fig. 3B**). We found that oncogenes amplified on circular amplicons achieved higher copy numbers than the same oncogenes amplified on Amplified-noncircular structures (**Extended Data Fig. 3C**). We further observed that the association between ecDNA structures and oncogene amplification did not extend to breakpoints. For 24 frequently amplified oncogenes, the frequency of observing a specific number of breakpoints in a unit interval decayed exponentially, consistent with random occurrence around the oncogene (**Fig. 1E**; **Extended Data Fig. 3D**, **Extended Data Fig. 3E**). These results suggest that ecDNA are formed through a random process, where selection for higher copies of growth promoting driver oncogenes leads to rapid oncogene amplification during cancer development and progression, in a way that also retains intratumoral genetic heterogeneity, due to its mechanism of uneven inheritance^3,16^.

Circular amplicons also differed from Amplified-noncircular amplifications in other notable ways. Circular and Amplified-noncircular amplifications showed similar likelihood of occurring in samples with chromosome-arm level aneuploidy (**Extended Data Fig. 4A**) and whole-genome duplication, which might arise as a result of chromosome missegregation ^17^ or other mitotic errors ^18^ (**Extended Data Fig. 4B**). Smaller and more focal genomic gains and losses result from different mutagenic processes, associated with genomic instability. We observed an increase in the number of DNA segments in samples marked by Circular amplicons, compared to other categories (**Fig. 2A**). The frequency of copy number losses was comparable between Circular and Amplified-noncircular amplicon samples (**Extended Data Fig. 4C**), but genomic segment gains were more frequently detected in samples with circular amplification (Wilcoxon rank sum test: p-val < 1e-14) (**Extended Data Fig. 4D**). This observation coincided with a threefold increase in gene fusion events inferred from matching RNAseq profiles (**Fig. 2B**; Binomial test: p-value <1e-138) compared to Non-circular amplification. Clustered mutations, also referered to as kataegis, were significantly more frequently detected in Circular amplicons relative to Amplified-noncircular amplicons, suggesting increased incidence of kataegis (Hypergeometric test: p-value≅ 0)(**Extended Data Fig. 4E**). The majority of Circular amplicon breakpoints showed no or minimal sequence homology (<5 bp), raising the possibility that non-homologous end joining could be involved in ecDNA formation. In contrast, Amplified-noncircular amplicon breakpoints showed significantly more micro-homologies than were seen on circular amplicons (**Extended Data Fig. 4F,** p-value<0.0005; two-sided Fisher’s exact test).

**Fig. 2.**
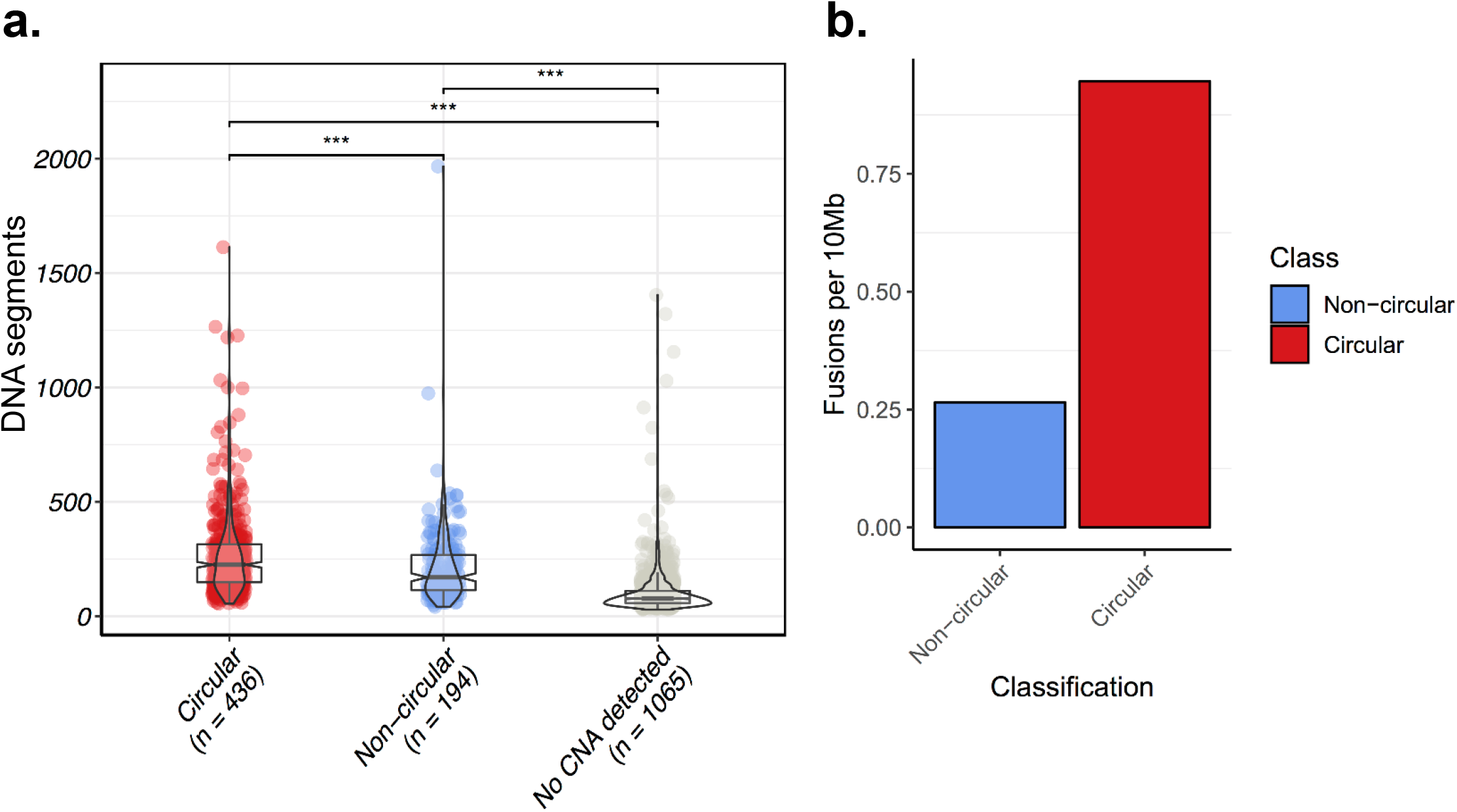
Total number of copy number segments and transcript fusions are increased in Circular amplicon tumor samples. **A.** TCGA copy number array data was used to count the total number of DNA segments within a sample. Circular samples contained statistically significantly more DNA segments than non-circular and no CNA detected (p-val < 1e-5 and 1e-128, respectively; Wilcox Rank Sum Test). **B.** Circular structures expressed significantly more gene fusions compared to non-circular amplicons, after size normalization.

We sought to examine the transcriptional consequences of circular ecDNA amplification at the population level. We detected a highly significant correlation between DNA copy number and gene expression level in all categories of DNA amplification, Circular and Non-circular. However, at comparable DNA copy number, oncogenes on Circular amplicons were significantly more highly expressed than those on Amplified-noncircular amplicons (p-value < 0.003; Wilcoxon rank sum test; **Fig. 3A**; **Extended Data Fig. 5**), showing a higher transcriptional rate (2.6X higher compared to Amplified-noncircular, 8.3X higher compared to oncogenes on non-amplified regions). To test if the epigenetic mechanisms governing gene expression were different between Circular amplifications and Amplified-noncircular regions, we analyzed the overlapping ATAC-seq profiles available for 24 samples^19^. The results (**Fig. 3B**) showed that chromatin of Circular amplicons was significantly more accessible compared to Amplified-noncircular categories (1.3 times higher ATAC-seq signal; Wilcoxon rank sum test; p-value < 0.003), suggesting that increased accessibility plays a role in dysregulation and higher expression of oncogenes on circular amplifications (ecDNAs).

**Fig. 3.**
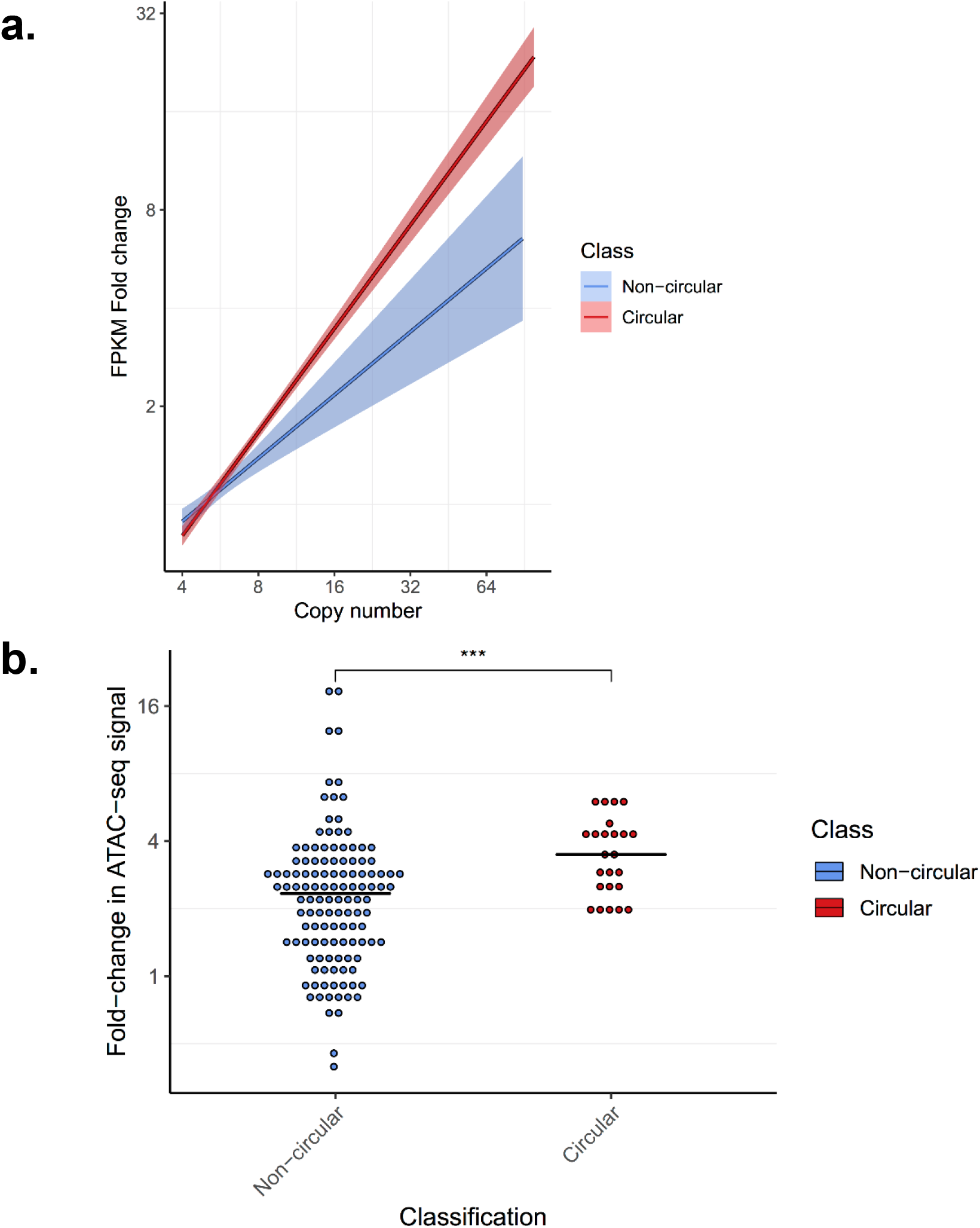
Gene expression and chromatin accessibility of amplicon classes. **A.** Copy number of the gene versus its fold-change in FPKM for all genes with a copy count greater than 4 and less than 100, for each gene on each amplicon. The fold-change in FPKM is computed as the gene’s (FPKM-UQ+1) divided by the average of (FPKM-UQ+1) for the same gene in all other tumor samples from the same cohort for which the gene is not on any amplicon (i.e., not amplified). Linear regression lines are shown for each classification class. Tukey’s range test shows genes on circular structures are significantly different to genes on non-circular structures (p-value < 1e-15). **B.** For each amplicon in the 24 TCGA samples with ATAC-seq and AmpliconArchitect results, the log2 fold-change in ATAC-seq signal across the amplicon relative to tissue types without amplification within the same region is shown. Each point represents a separate amplicon. The distribution of fold-change for circular amplicons is statistically significantly higher than non-circular (Wilcoxon rank sum test; p-value < 0.003).

Having developed a way to stratify tumors based upon amplification architecture, we examined the impact of ecDNA-based amplification on two hallmarks of cancer, immune evasion and cell proliferation. We used previously developed gene expression signatures^20^ to evaluate the distribution of immune infiltrate and cell proliferation scores by amplicon grouping. The cellular proliferation but not immune infiltration pathway scores were significantly higher (**Fig. 4A**, p-val < 1e-7; Wilcoxon Rank Sum Test; **Extended Data Fig. 6**) in the Circular amplification category compared to the other two groups. We did not observe difference in activity of the immune signature score between groups (**Fig. 4A**, p-val < 0.03; Wilcoxon Rank Sum Test). The increased activity of the cell proliferation gene signature suggested a higher rate of proliferation and tumors that behave more aggressively.

**Fig. 4.**
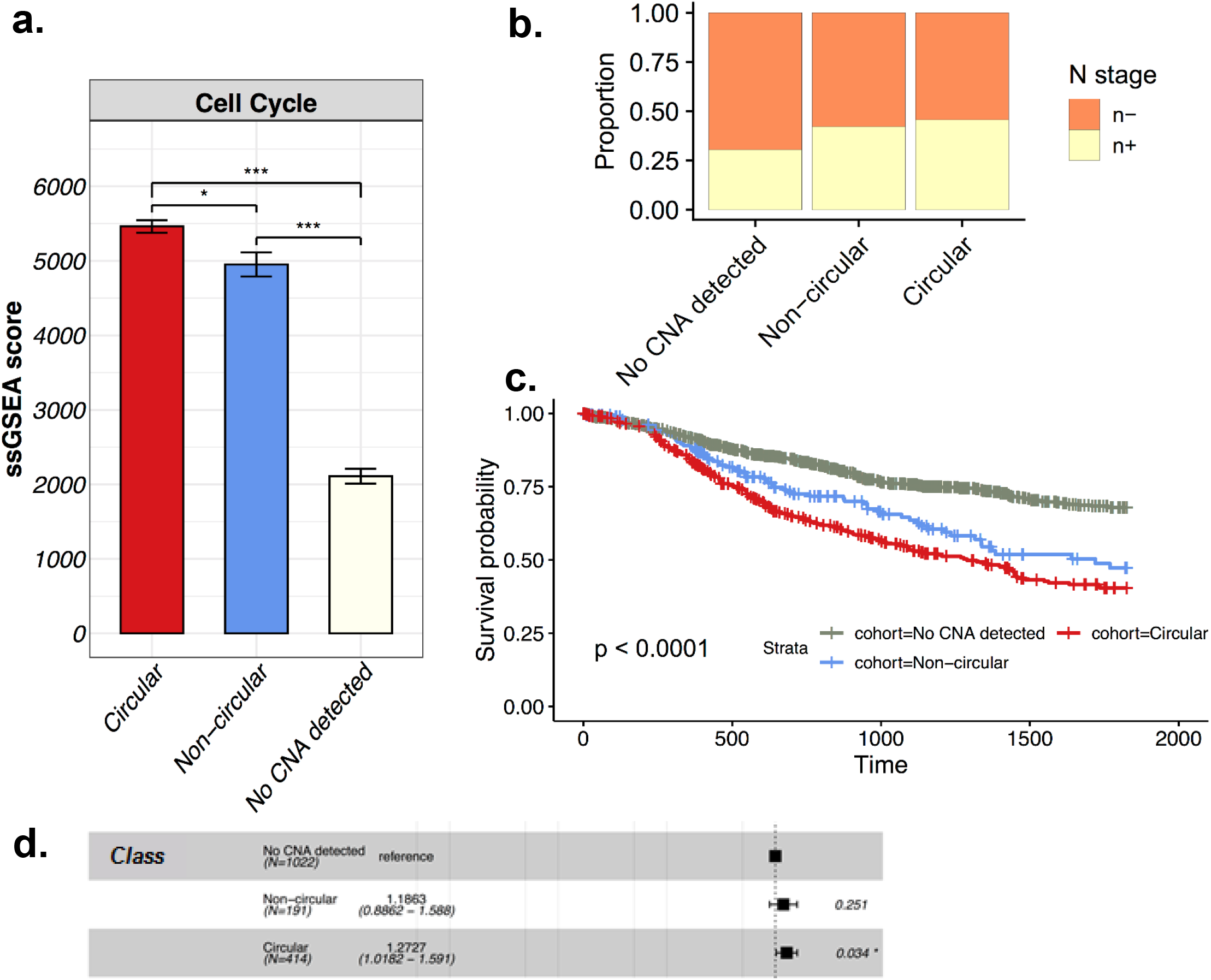
Presence of circular amplification associates with poor outcomes. **A.** Cell proliferation gene expression signature single sample GSEA (ssGSEA) scores by amplification category. Shown are means and 95% confidence intervals of the ssGSEA scores. Samples with circular structures showed significantly higher ssGSEA scores than samples with non-circular amplicons. **B.** Lymph node stage for primary tumors showing samples with amplification are more likely to have spread to the lymph node at time of diagnosis. **C.** Kaplan-Meier five-year survival curves by amplification category. Both amplified-noncircular and circular amplification have significantly worst outcome than No CNA samples (p-val < 1e-4 and 1e-15, respectively). Circular amplification has worse but not significant outcome compared to amplified-noncircular (p-val < 0.07). **D.** Cox-Hazard model, incorporating disease and patient cohorts as parameters showing circular amplification results in significantly higher hazard rates.

To determine whether cancers that have ecDNA amplification were associated with tumor progression, we examined the impact of circular amplification on lymph node status at initial presentation, and overall survival. We found that the proportion of cases in which the tumor had spread to a lymph node at the initial time of diagnosis was significantly increased in tumor samples that had either circular or non-circular amplification (**Fig. 4B**; p-value < 0.02 no- CNA vs Amplified-noncircular, p-value < 1.0e-05 Circular-amplicon vs Amplified-noncircular). Additionally, we found a significant difference in overall survival of patients stratified by amplification category. Patients whose tumors contained circular amplification associated with significantly worse overall outcomes compared to patients whose tumors harbored either non-circular amplifications or no amplifications (**Fig. 4C**; p-val < 1e-15 versus no-CNA detected; p-val < 0.07 against Amplified-noncircular; Log-rank test). To account for the possibility that differences in survival rate are being influenced by the disease subtype, as circular amplicons are much more prevalent in aggressive cancers such as glioblastoma, we fit the data to a Cox Hazard model that tested survival after controlling for disease subtype. The model showed that patients with circular amplicons had significantly higher hazard rates (**Fig. 4D**; 28% increase in hazard rate relative to no-CNA, p-val < 0.03).

Cancer genomics is itself evolving from reading out the “code” to unraveling its function. The 3D organization of the genome plays a critical role in determining how that genome functions, or malfunctions, as occurs in cancer. The data presented here demonstrate that ecDNA play a critical role in cancer, providing a mechanism for achieving and maintaining high copy oncogene amplification and diversity. This mechanism of amplification is operant in a large fraction of human cancers, and contributes to the poor outcomes for patients. The potential to leverage the presence of ecDNAs in a quarter of human cancers for diagnostics or therapeutics provides a link between cancer genomics and broad utility for patient populations

## METHODS

### AmpliconArchitect

We used AmpliconArchitect^11^ infer the architecture of the ‘amplicons’ --- large (>10kb) rearrangements with high copy numbers (CN>4) that are inferred to have co-amplified as a structure. AmpliconArchitect takes as an input aligned WGS sequences and seed intervals of the amplicon. AmpliconArchitect then searches for other regions that belong to the amplicon by exploring the seed intervals, and extends beyond the intervals if it encounters copy number changes or discordant edges that support a breakpoint. The collection of intervals and breakpoints are combined to form a fine network with nodes representing segments and edges representing rearrangements, which we call the breakpoint graph. This breakpoint graph is can be further decomposed into simple cycles to identify any circular paths within the amplicon structure, which is indicative of ecDNA presence. The detected amplicons were annoted with the Ensembl Release 75 gene database (GRCh37).

### Amplicon and sample classification

As a perquisite, amplicons must contain ≥ 10kb of genomic segments amplified to at least four copies above median ploidy in order to be considered a valid amplicon. We then use the AmpliconArchitect derived breakpoint graph to classify amplicons into three categories: 1. Circular amplification; 2. Heavily rearranged amplification; and, 3. Amplified-noncircular (**Extended Data Fig. 1A)**. Amplicons were denoted as Circular amplification if the segments form a cycle in the graph of total size at least 10kb and has at least a copy count of four. Non-circular amplicons were denoted as heavily rearranged if the breakpoints connect segments from different chromosomes, or distal (>1Mb) regions (**Extended Data Fig. 1A)**. Non-circular, non-distal amplicons were denoted as locally rearranged. All other regions that were not part of any amplicon structure were classified as not-amplified. While an amplicon may fit the requirements for several categories (i.e., a circular amplicon may also comprise heavily rearranged amplifications), priority was given to the circular amplification category, followed by heavily rearranged and finally amplified-noncircular. Similarly, samples were classified based upon what amplicons are present within the sample, giving precedence to the presence of amplicon with highest priority. For example, a sample with both circular and heavily rearranged amplification would be classified as circular amplification. Samples without any amplicons are classified as ‘No CNA detected’.

### Cell line validation

We ran AmpliconArchitect on the whole-genome sequencing data from 11 glioblastoma neurosphere cultures from deCarvalho et al ^1^. with FISH images and the 23 cell lines from different cancer types from the Turner et al ^2^ study. Fluorescence in-situ hybridization (FISH) was performed in parallel, as described. The seed interval for each cell line included the probe region. For each probe, we reported whether it landed in an amplicon (inferred from AmpliconArchitect), and if so what was the amplicon classification. The distribution of the average ecDNA per cell was computed as the average number FISH probes that co-localized on ecDNA across all the images for that particular cell line+FISH probe combination (**Extended Data Fig. 1B**). Wilcoxon Rank Sum test was used to detect significant differences in average ecDNA counts per cell across the amplicon classes.

### TCGA processing

We processed TCGA whole genome sequencing BAMs through the Institute for Systems Biology Cancer Genomics Cloud (https://isb-cgc.appspot.com/) that provides a cloud-based platform for TCGA data analysis. We used genome-wide snp6 copy number segments with copy number log ratio equal to 1 as seed interval(s) of interest that are required for the input to AmpliconArchitect^11^. Default parameters and reference files were used for all other settings. Details on how to run AmpliconArchitect have been described in the corresponding manuscript^11^ and its source code depository.

We ran AmpliconArchitect on tumor and normal WGS samples from 1979 patients. Samples were classified based upon the amplicon with highest precedence present in the sample, or classified as ‘No CNA detected’ if no amplicons are present in the sample. Samples classified as highly-rearranged are removed from further analysis due to the ambiguity of ecDNA status of the sample.

### TCGA processed data

The processed data (hg19) and clinical data were found at the GDC (https://portal.gdc.cancer.gov/legacy-archive/search/f) and the PancanAtlas publication page (https://gdc.cancer.gov/about-data/publications/pancanatlas). Somatic muations for TCGA whole genome sequencing were downloaded from the ICGC PCAWG portal (https://dcc.icgc.org/pcawg)^21^.

### Oncogene analysis

We examined the enrichment of the 24 recurrent oncogenes known to be activated by amplification by counting the total number of base pairs from the amplicon classes from all the tumor samples that overlap these oncogenes. We then simulated 10,000 replicates by sampling random regions of the same size of the amplicons and computed an empirical expected distribution of base pairs covering the oncogenes if the amplicons were randomly sampled across the genome. We report the z-score between the empirical distribution and observed value for the amplicon classes. We also report the average copy count, estimated from AmpliconArchitect. For each of these oncogenes on an amplicon structure, we reported the position of breakpoint detected within a 1 MB region flanking the oncogene using the breakpoint graph to infer breakpoints. We partitioned the region into 1000 bp windows and counted the total number of breakpoints that landed in each window, and display a histogram of these counts. We modeled the histograms using an exponential distribution and show that under the assumption that the breakpoints are distributed randomly, the histograms closely follow the exponential distribution.

We used *allOnco* (http://www.bushmanlab.org/links/genelists), a set of 2,579 cancer genes generated from curated collections cancer genes from many different publications. We identified all amplicons that overlapped with the oncogenes and report the proportion amplified oncogenes that are circular.

### Genomic instability analyses

We computed total copy number gains/losses as the number of snp6 copy number segments with copy number >=4 or <= 1. Wilcoxon Rank Sum test was used to test for a significant difference between the two distributions. We used the data from a previous study^22^ to estimate the genome doubling status and chromosomal arm duplication and loss for each sample. Wilcoxon Rank Sum test was used to test significance between the distribution of gains and losses and Chi-squared test was used to test significance between the distribution of whole genome doublings.

We used the data from the TCGA fusion database (https://tumorfusions.org/)^23^ to identify fusions events that occur on an amplicon. For each fusion in the database, we consider it valid if both ends of the fusion breakpoint junction occur on the same amplicon. In total, 710 amplified fusions were detected. We computed the average fusion events per 10 Mb as the total number of fusions that landed within an amplicon class divided by the sum of all the base pairs of the amplicon class multiplied by 10e7. To test whether circular amplicons were enriched fusion events, we computed the p-value of observing at least the number of fusion events on circular amplicon under a binomial distribution where the probability ***p*** was estimated using the total number of fusion events on the amplified-noncircular divided by the total base pairs of the amplified non-circular event and the number of trials ***n*** as the total base pairs of the circular amplicons.

### RNAseq and ATACseq analyses

Of the 1,695 tumor samples, 1,608 had RNA-seq data in the format of FPKM-UQ expression data. For each gene within each disease cohort, we computed a baseline FPKM-UQ as the average FPKM-UQ of all samples for which the gene was not found on an amplicon (i.e., average expression of the unamplified gene). We then computed the fold-change in expression of each gene on each amplicon as the FPKM-UQ of the amplified gene divided by the average FPKM-UQ of the unamplified samples, and report the distribution of fold-changes versus the copy number. Tukey’s range test was used to test significance between slope of the FPKMs for circular and amplified-noncircular.

ATAC-seq profiles were available for 24 samples. For each amplicon in each sample, fold-change in ATAC-seq signal was computed as the average ATAC-seq signal across the amplicon region divided by the average ATAC-seq signal for the same region in the unamplified samples of the same cancer type. Wilcoxon Rank Sum test was used to test significance between the two distributions.

### Kataegis

Localized mutation clusters (kataegis loci) were defined as having 6 or more consecutive mutations with an inter-mutation distance of < 1kb in a similar way to a previously used approach ^24^.

### Inferring breakpoint homologies

For each breakpoint, sequencing reads around +/− 1000 bps of the breakpoint were locally reassembled with SvABA^25^ to produce a contiguous consensus sequence of each breakpoint, precise breakpoint positions, and the level of homology at breakpoints.

### Statistical analysis

Survival curves were estimated with the Kaplan–Meier method, and comparison of survival curves between groups was performed with the log-rank test in R survival package. Hazard ratios were estimated with the Cox proportional hazards regression model in the survival R package.

## ACKNOWLEDGEMENTS

This work was supported by the Ludwig Institute for Cancer Research (P.S.M.), Defeat GBM Program of the National Brain Tumor Society (P.S.M.), NVIDIA Foundation, Compute for the Cure (P.S.M.), The Ben and Catherine Ivy Foundation (P.S.M.), generous donations from the Ziering Family Foundation in memory of Sigi Ziering (P.S.M.), and Ruth L. Kirschstein National Research Service Award This work was also supported by the following NIH grants: NS73831 (P.S.M.), GM114362 (V.B.), R01 CA190121 and Cancer Center Support Grant P30 CA034196 (R.G.W.V), R35CA209919 (H.Y.C.), RM1-HG007735 (H.Y.C.), and NSF grants: NSF-IIS-1318386 and NSF-DBI-1458557 (V.B.); and grants from the Musella Foundation and the B*CURED Foundation (R.G.W.V). H.Y.C. is an Investigator of the Howard Hughes Medical Institute. The results published here are in whole or part based upon data generated by the TCGA Research Network: https://www.cancer.gov/tcga. Analysis of TCGA was made possible through the Cancer Genomics Cloud of the Institute for Systems Biology (ISB-CGC).

---

**Extended Data Fig. 1.**
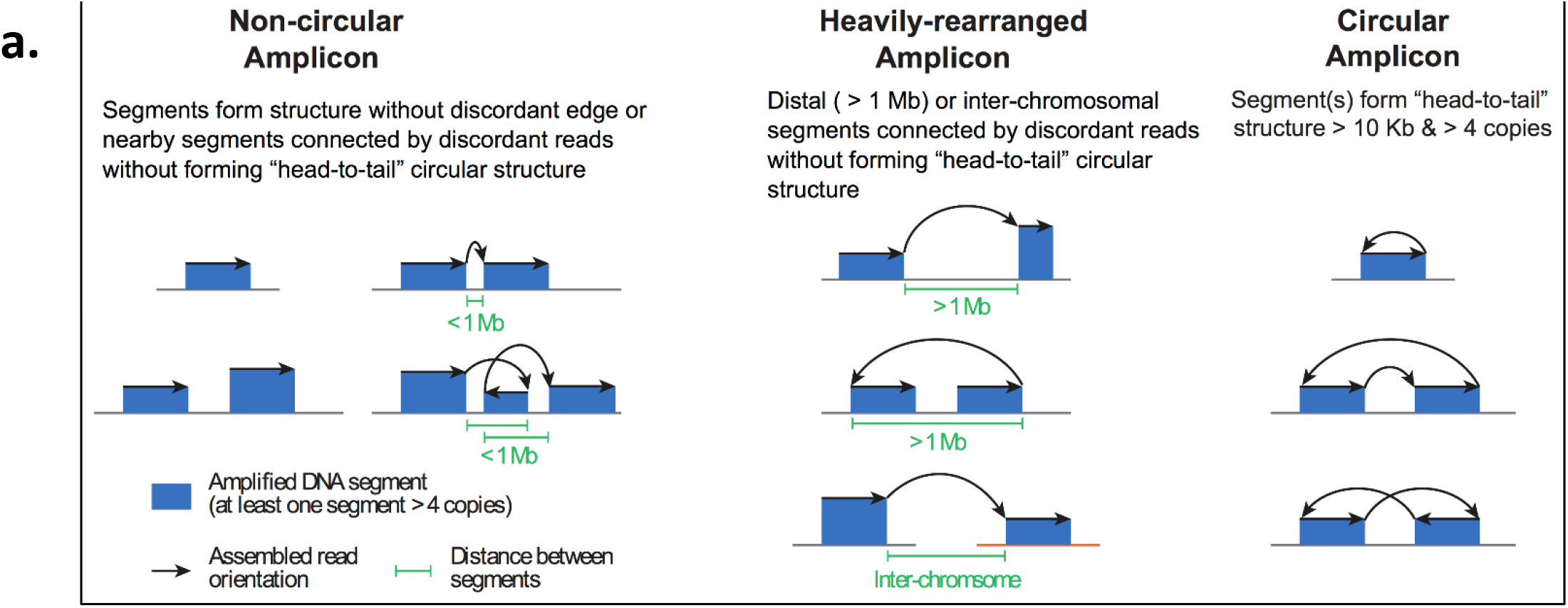
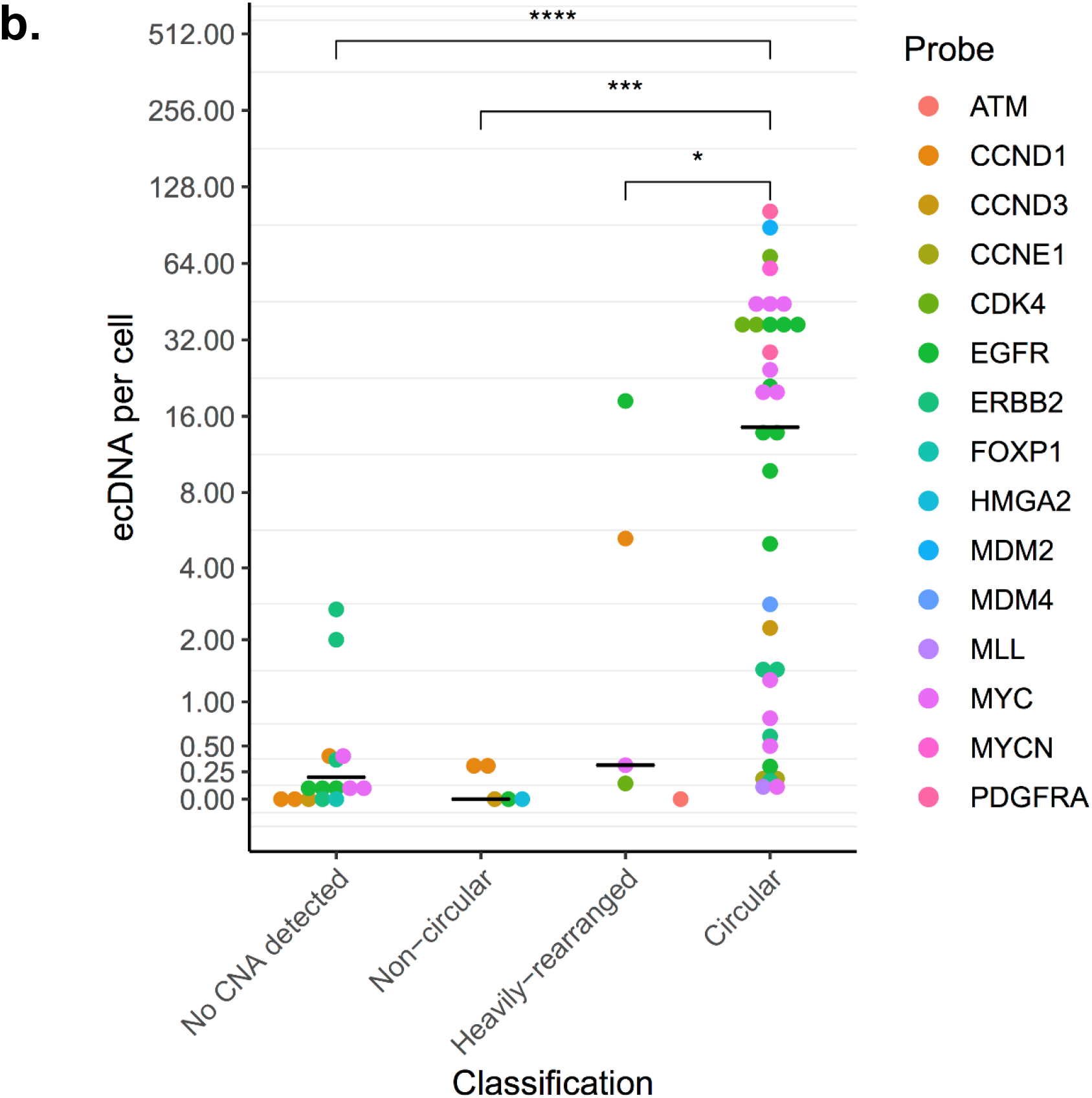
Amplicon classification. **A**. Schematic representation of the three classification categories. Amplicons are classified using a hierarchical scheme based upon the genomic reconstruction of the amplified regions (i.e., any region with a copy count of 4 or greater) and the presence or absence of discordant edges between these regions. Amplicons must have a minimum 10kb of amplified regions in order to be considered a valid amplicon. The first category is circular amplicon, which is an amplicon that contains one or more amplified segments forming a cyclic path of at least 10kb bps in length and has an average amplification of four copies. The second category is heavily-rearranged amplicon, which is an amplicon that contains amplified segments that are connected by discordant read pairs, and at least one breakpoint junctions is inter-chromosomal or greater than 1Mb is size. The third category is non-circular amplicon, which is any amplicon that contains amplified segments with no discordant edges or with discordant edges, but all breakpoint junctions are less than 1 Mb in size. All other regions are considered not amplified. As the classification scheme is hierarchical, each amplicon can only have one class, and the highest rank class has precedent (i.e., an amplicon that is both circular and heavily-rearranged will be classified only as circular). As samples can have multiple amplicons, the sample is classified as the amplicon with highest precedent (i.e., a sample with 1 circular amplicon and 3 heavily-rearranged amplicons would be classified as circular). **B**. Validation on cell line data. Validation of the classification scheme on cell line data with FISH experiments for detecting ecDNA from the Turner et al. and deCarvalho et al. studies. FISH probes were designed for selected oncogenes and DAPI staining was performed to determine whether the FISH probe landed on chromosomal DNA or ecDNA. For each cell (represented as an image of the cell in metaphase), the number of positive ecDNA probes were counted, and for each cell line, the average positive ecDNA per cell was reported. For each probe, we report whether it landed in an amplicon (inferred from AA), and if so, what was the amplicon’s classification. The distribution for the average ecDNA per cell between the circular and heavily rearranged (p-value < 0.05; Wilcoxon rank sum test) and No CNA detected/heavily rearranged were statistically significantly different (p-value < 0.001; Wilcoxon rank sum test).

**Extended Data Fig. 2.**
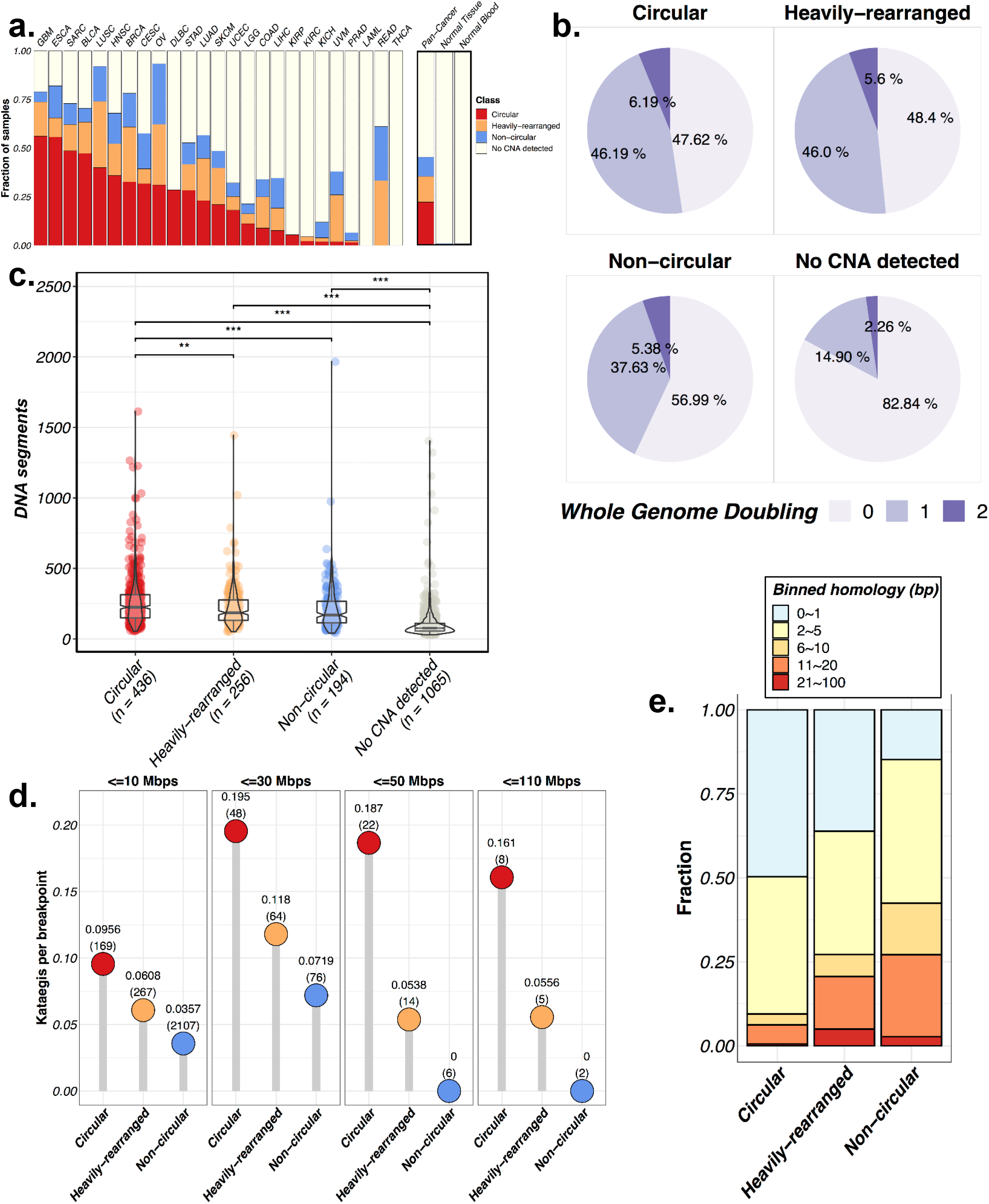
**A.** summary result including heavily-rearranged category**. B.** Genome doubling events by amplification class. **C.** Total number of copy number segments by amplification class. **D**. Kataegis frequency differences between amplification categories. **E.** Breakpoint homology by amplification class.

**Extended Data Fig. 3.**
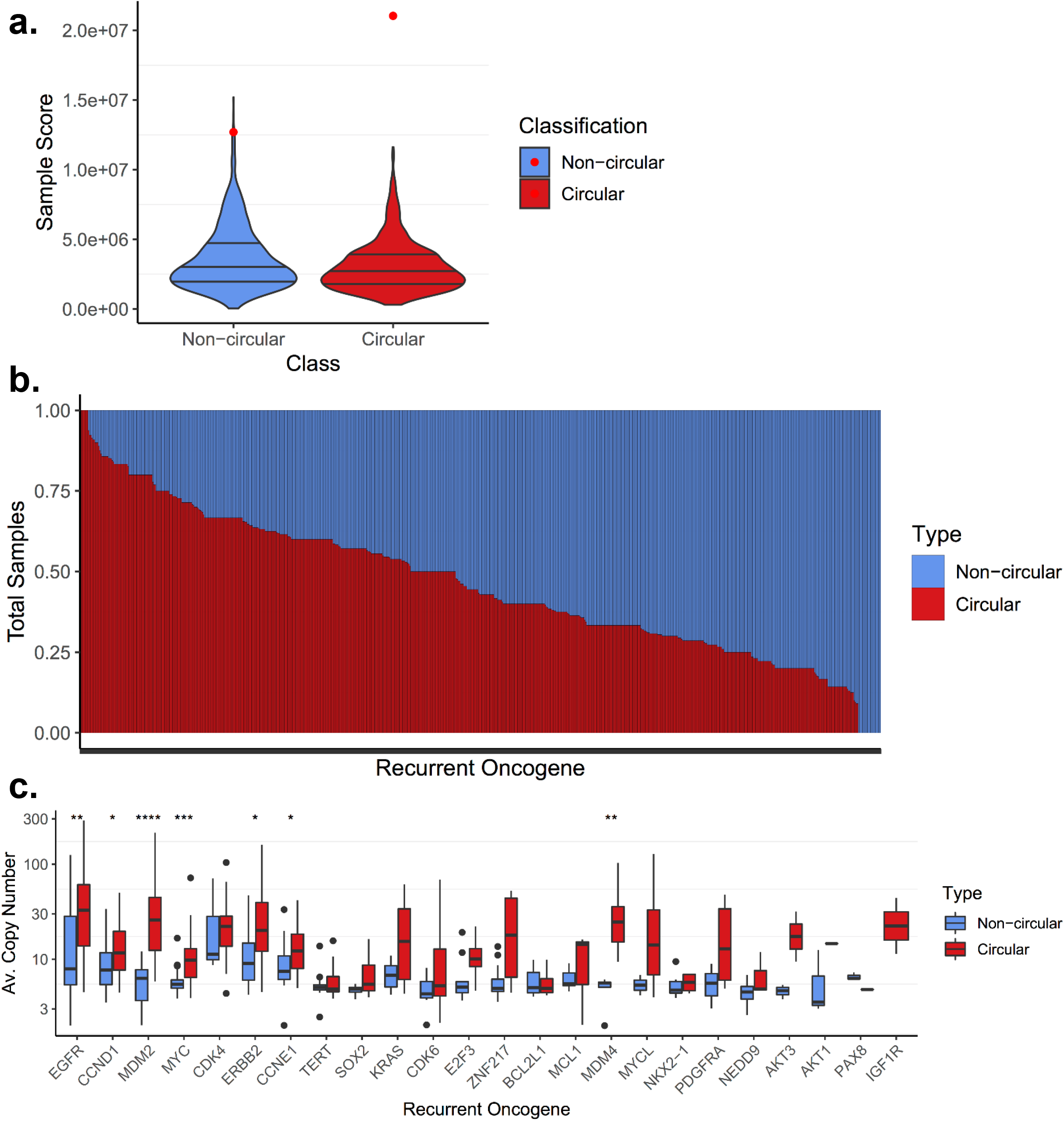
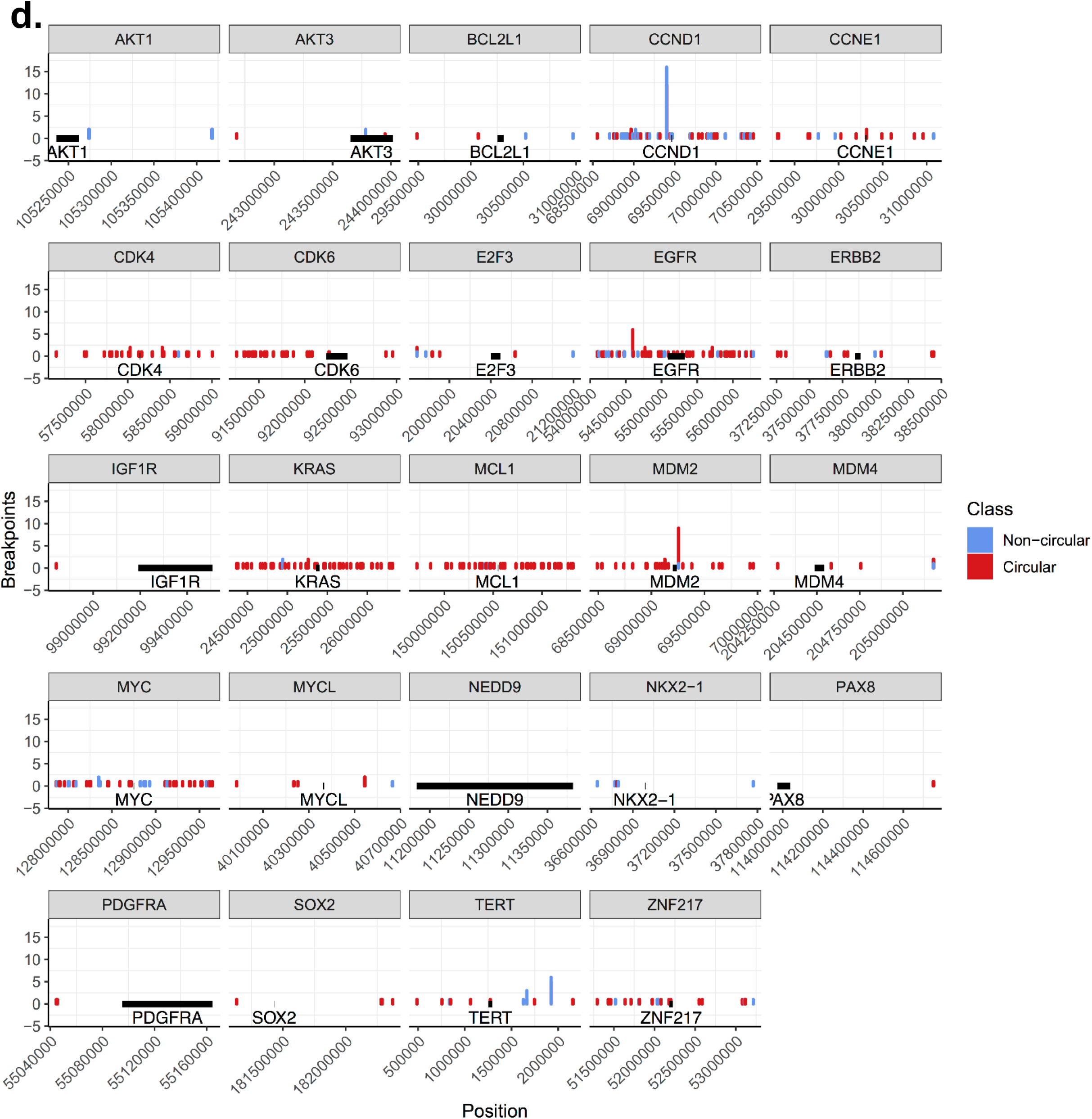
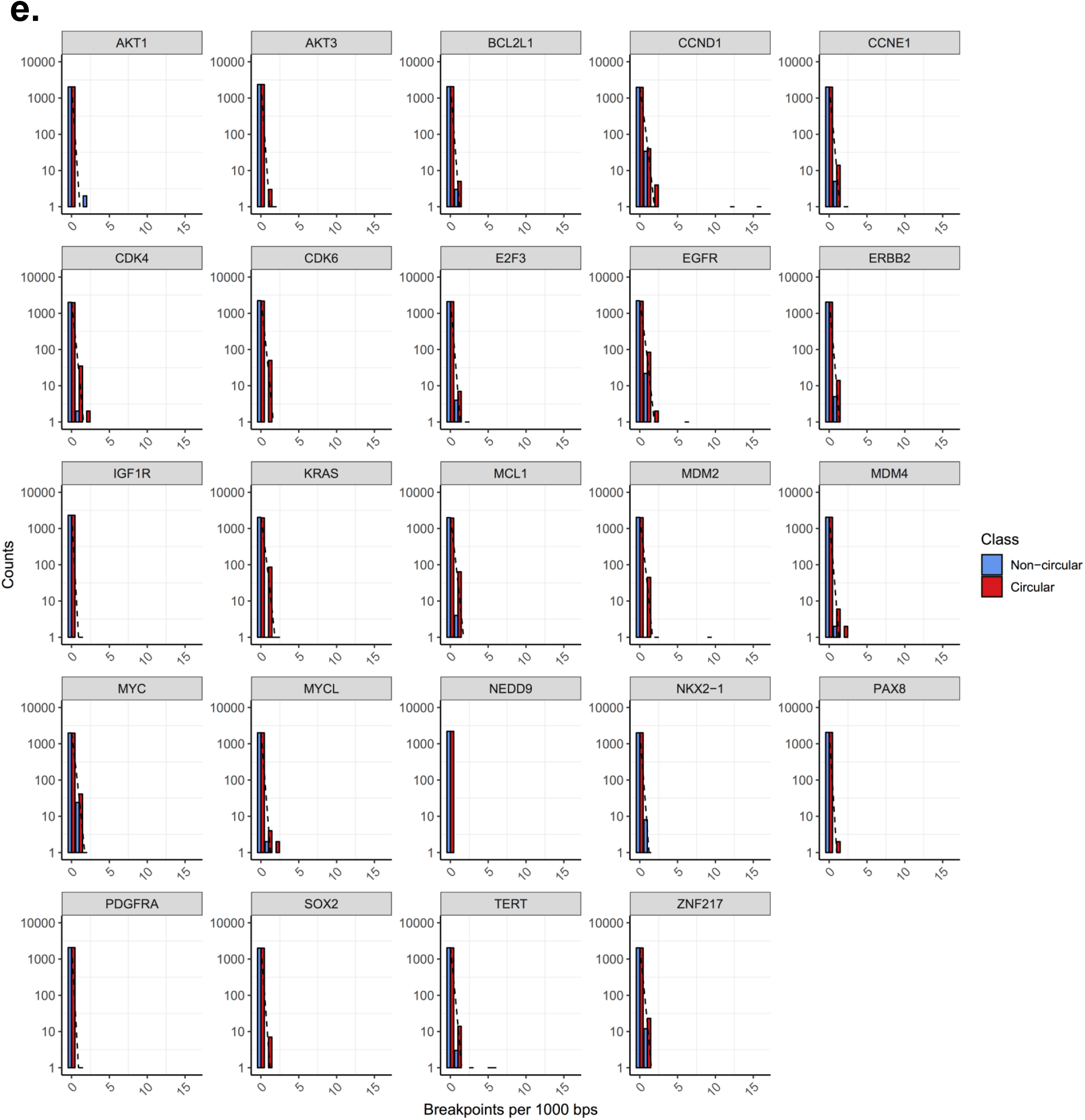
Circular vs amplified non-circular amplification comparisons. **A**. 24 recurrently amplified oncogenes significantly overlap circular regions (z-score 10.9), especially compared to amplified non-circular (z-score 4.0). **B**. For all oncogenes with copy number >= 4 (defined from the DNA copy number array data) and present in at least 5 samples, we show the class distribution of that oncogene. The oncogenes are ordered by proportion on circular amplification. **C**. For the 24 recurrent oncogenes known to be activated via amplification (**Zack et al. Nat Gen. 2013**), we report the average copy number for the oncogenes for circular amplification versus amplified-noncircular amplification. **D.** Breakpoint locations across the 24 recurrent oncogenes activated by amplification. Outliers in CCND1 and MDM2 were results of mapping bias due to short ALU repeats near the oncogene region. **E.** Breakpoint locations across the 24 recurrent oncogenes activated by amplification.

**Extended Data Fig. 4.**
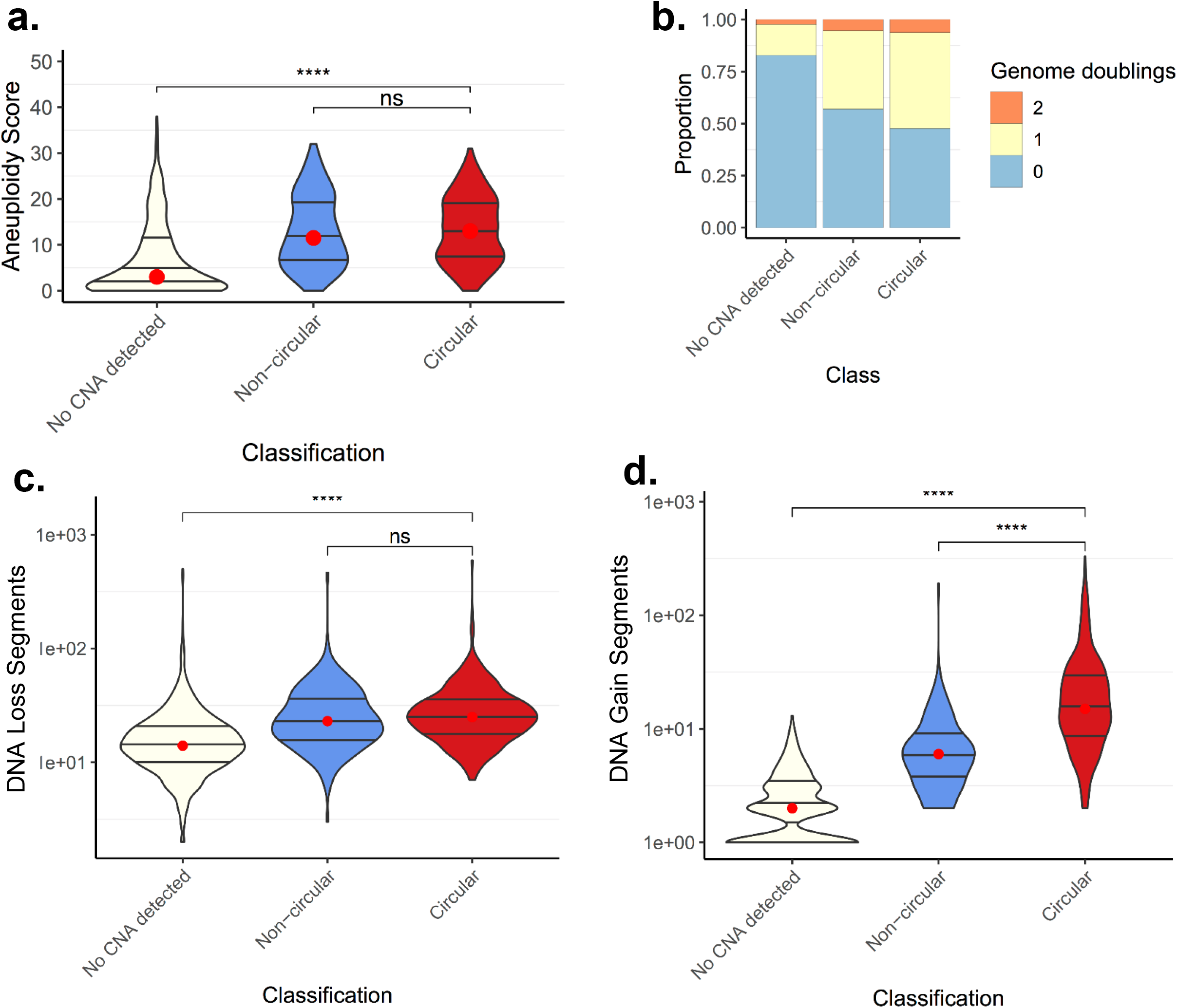
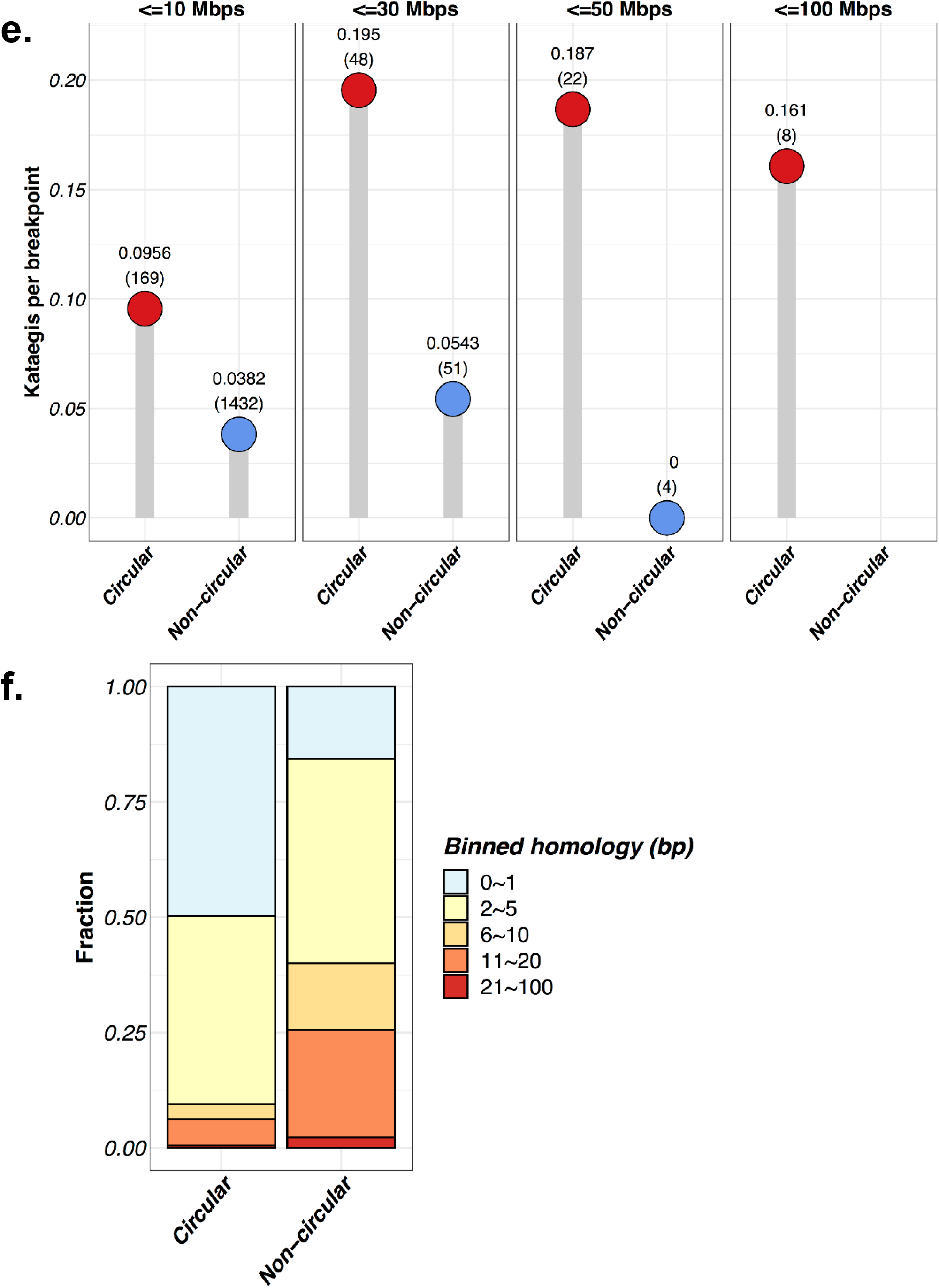
Aneuploidy and genomic instability events by amplification class. **A.** Chromosome arm aneuploidy scores showing no difference in chromosomal arm level events between amplified-noncircular and circular amplification. **B.** Genome doublings distribution across classes showing no difference in distribution between amplified-noncircular and circular amplification. Circular amplification and non-amplified are different (Chi-square test; p-val < 1e-12). Circular amplification and amplified-noncircular are not different (Chi-square test; p-val < 0.10). **C.** Distribution for total DNA loss segments by amplification class. TCGA CNV array data was used to count the total number of DNA losses within a sample. A DNA loss was defined as a segment with CN <= 1. **D.** Same as C, but for gain segments (CN >=4). Circular samples contain statistically significantly more DNA gains than non-circular and no-CNA detected (p-val < 1e-14 and 1e-127, respectively; Wilcox Rank Sum Test). Non-circular contain statistically significantly more DNA gains than no-CNA detected (p-val < 1e-35). **E.** Kataegis frequency differences between amplification categories. Amplicons were grouped into Amplicon-size bins, and # kataegis frequency was normalized for the number of DNA breakpoints, demonstrating a higher occurrence of kataegis in Circular compared to Non-circular amplifications. The number of amplicons used is shown in parentheses. **F.** Breakpoint homology by amplification class. Note that inserted sequences were excluded.

**Extended Data Fig. 5.**
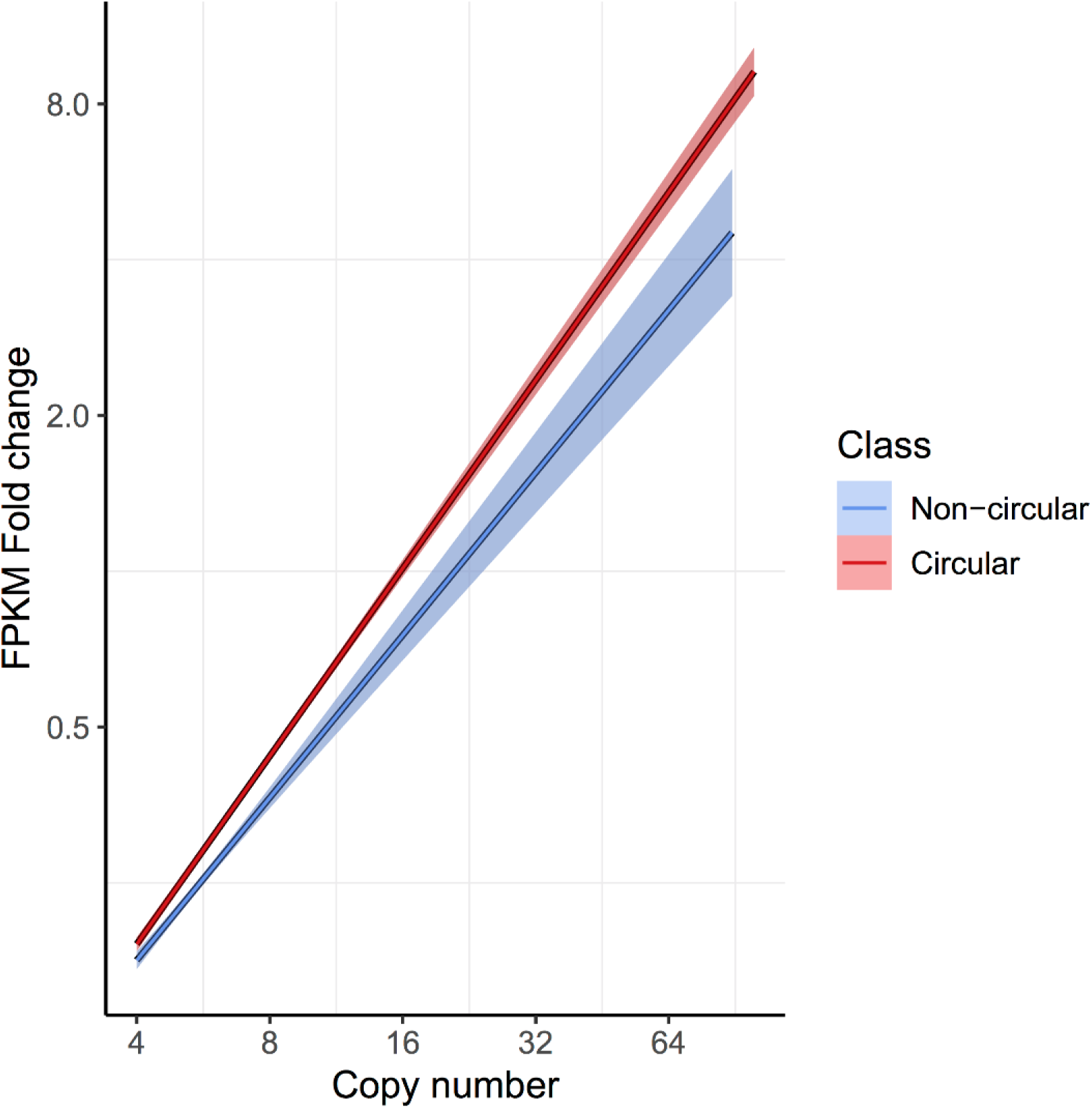
FPKM fold-change versus copy number. For each gene on each amplicon, we report the copy number of the gene versus its fold-change in FPKM for all genes with a copy count greater than 4 and less than 100. The fold-change in FPKM is computed as the gene’s (FPKM-UQ+1) divided by the average of (FPKM-UQ+1) for the same gene in all other tumor samples from the same cohort for which the gene is not on any amplicon (i.e., is not amplified). Linear regression lines are shown for each classification class. Tukey’s range test shows genes on circular structures are significantly different to genes on non-circular structures (p-value < 1e-15).

**Extended Data Fig. 6.**
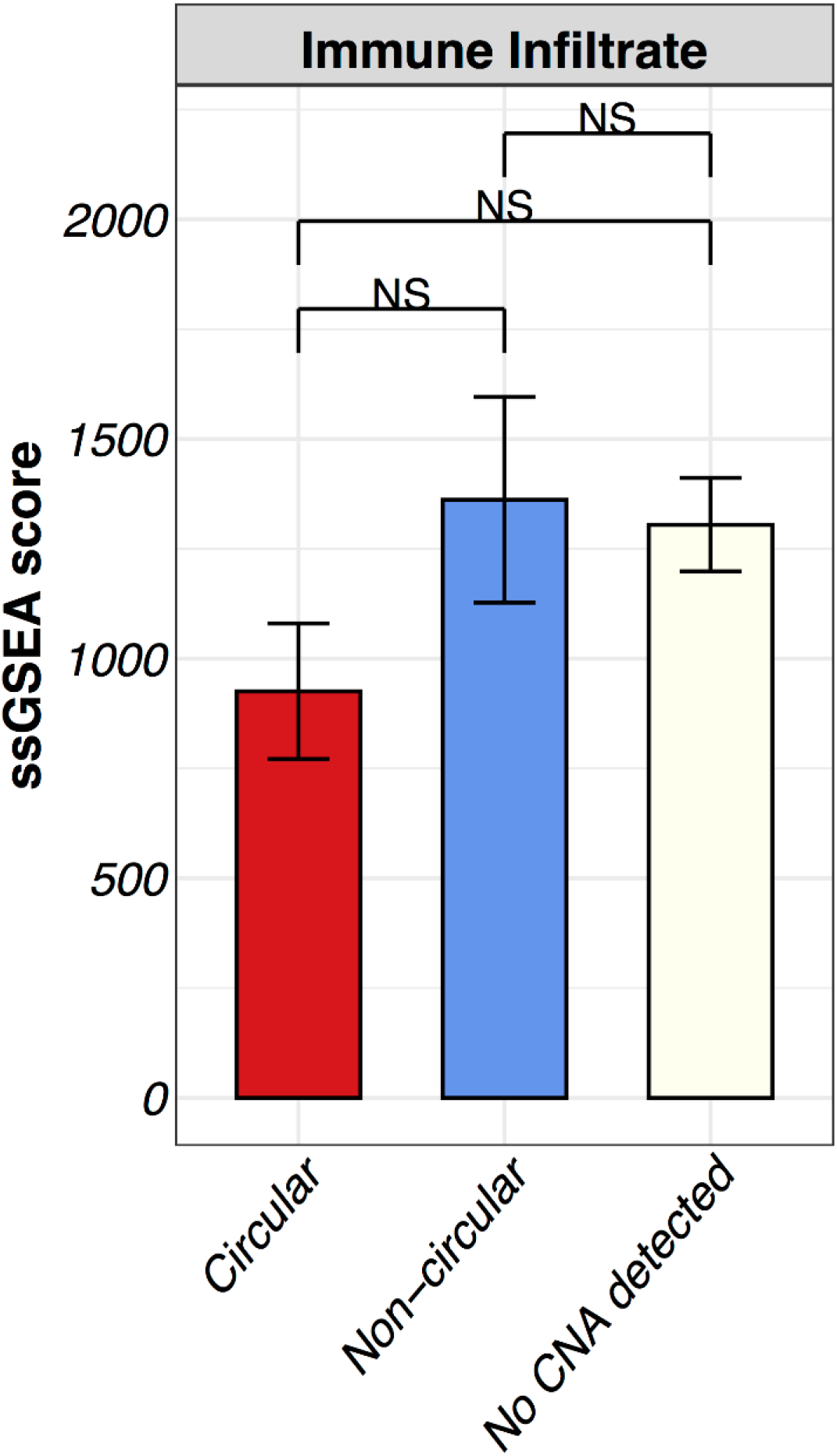
Circular amplification associates with worse outcomes. Immune gene expression signature single sample GSEA (ssGSEA) scores by amplification category. Shown are means and 95% confidence intervals of the ssGSEA scores. No significant difference was observed between classes.

